## Supplementary material for "AlphaFold2 modeling and molecular dynamics simulations of an intrinsically disordered protein": Supplimentary Figures 1-6

***Contents:***

**The amino acid sequence of *Nvjp-1***

**Figure S1**. Circular representation of the residue interaction networks.

**Figure S2**. Residue interaction networks from 100 ns MD trajectories of AF2 models.

**Figure S3**. Comparison of the single chains in 25 *Nvjp-1* homodimers

**Figure S4**. Residue interaction network from the MD trajectory of *Nvjp-1* dimer models C.2 and D.0.

**Figure S5**. Percentages of secondary structure elements (SSEs) estimated in 1000 AF2 models

**Figure S6**. The RGN2 model of *Nvjp-1* monomer.

### The amino acid sequence of *Nvjp-1*

>Nvjp-1

```
HNDGYGHDDHHGHGHGGYGGHGHGDYGGHGHGGYGGHGHGHGHGHFDD
HPFYTIPAFGHGYGHGHGGHGHGYGGHDGYGGHGGYGGHGGYGDHGHGGH
GYGSHGGHGQDYGGDYGGHGHGGHHHGGHDHDDFGHDFGHHGGDHGHHGG
GHHGHHDGYGHDQGHGHGHGDYDYSHGNQHDNGHREHYGGHQAAGHHGHG
ESGSHYGGQGGGGGSHHSGGHDGSHNHGKFQTQYSYRGNDKYGGHNQYQG
HGAYNEYSKSKGTGKYVAHGTEEGHHQLDGYEKDNHQGKYEGHGEHHSHQ
HGDGHHKRKEHGDHAGISGFHGQGHVQGNHALHGHHGGGHGHHGGEHGHS
ARGHHGGGGYGGGGHGHRGHHGGHQGHHGHH
```

Note: The signal peptide is not included (381 AA’s). The Gly (green) and aromatic (His, Tyr and Phe, black bold) residues are highlighted.

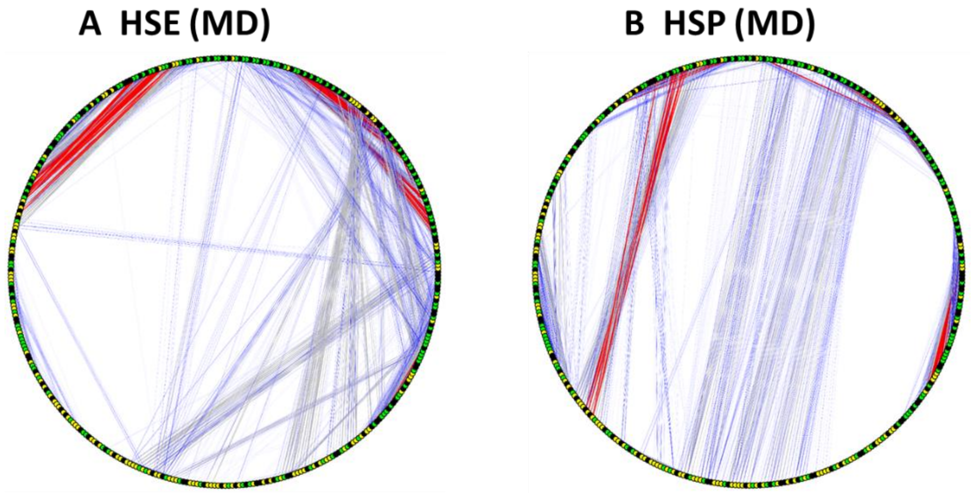


**Figure S1**. Circular representation of the residue interaction networks (also see Fig3D/E in the main text. The edge weights are averaged from MD simulations (**A**) in HSE state and (**B**) in HSP state, respectively. Interactions appear in >50% of the snapshots during MD are shown in red, the edges appear in <10% of the MD trajectory are shown in blue, and those in between are in grey. The green nodes represent Gly, black nodes are aromatic (H, Y and F) residues in *Nvjp-1*. All other nodes are colored in yellow.

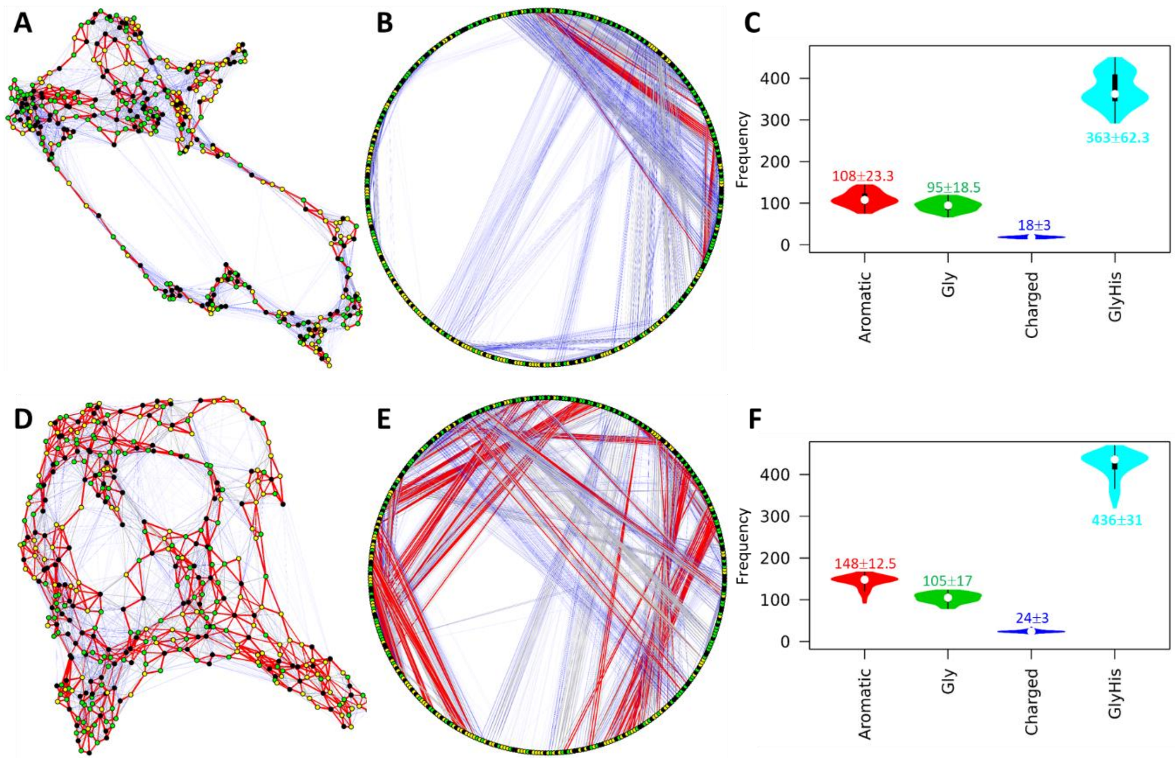


**Figure S2**. Residue interaction networks from 100 ns MD trajectories of AF2 models of C.2 (**A-C**) and D.3 (**D-F**) of *Nvjp-1* monomer. Fig 1B in main text compared the RMSDs of all monomer models.

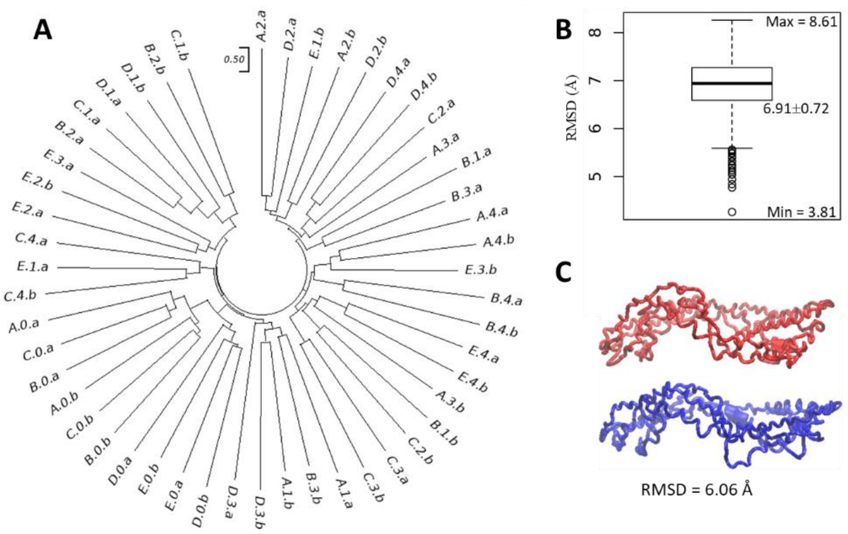


**Figure S3**. Comparison of the single chains in 25 *Nvjp-1* homodimers. (**A**) A structure-based phylogenetic tree. The letters "a" and "b" refers to two different chains in the same homodimer, respectively. (**B**) A box plot of the RMSD matrix. (**C**) Two chains (blue and red) from one of the models (A.0) has a $C_\alpha$ RMSD of 6.06 Å.

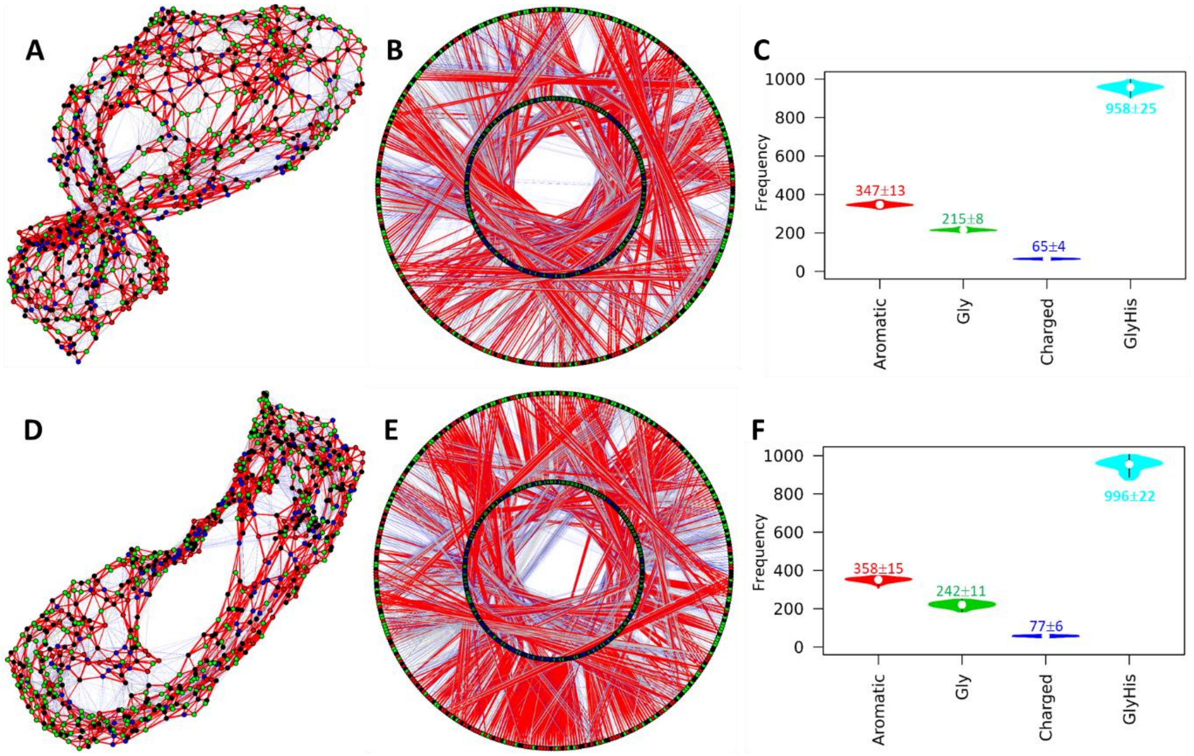


**Figure S4**. Residue interaction network from MD trajectories of *Nvjp-1* dimer models C.2 (**A-C**) and D.0 (**D-F**). Gly and His residues are in green and black, and other residues from chain A and B are in blue and red, respectively. Randomized (**A, D**) and circular (**B, E**) representations of the RINs are shown. The inner circle of the circular network is for chain A and the outer circle is for chain B, respectively. The interactions appear in >50% of all configurations from MD are in red edges (strong interactions) and the interactions appear in <10% are shown in blue edges (transient interactions), and those in between are shown in grey edges. Violin plots of interaction frequencies of selected reside groups are shown in **C** and **F**.

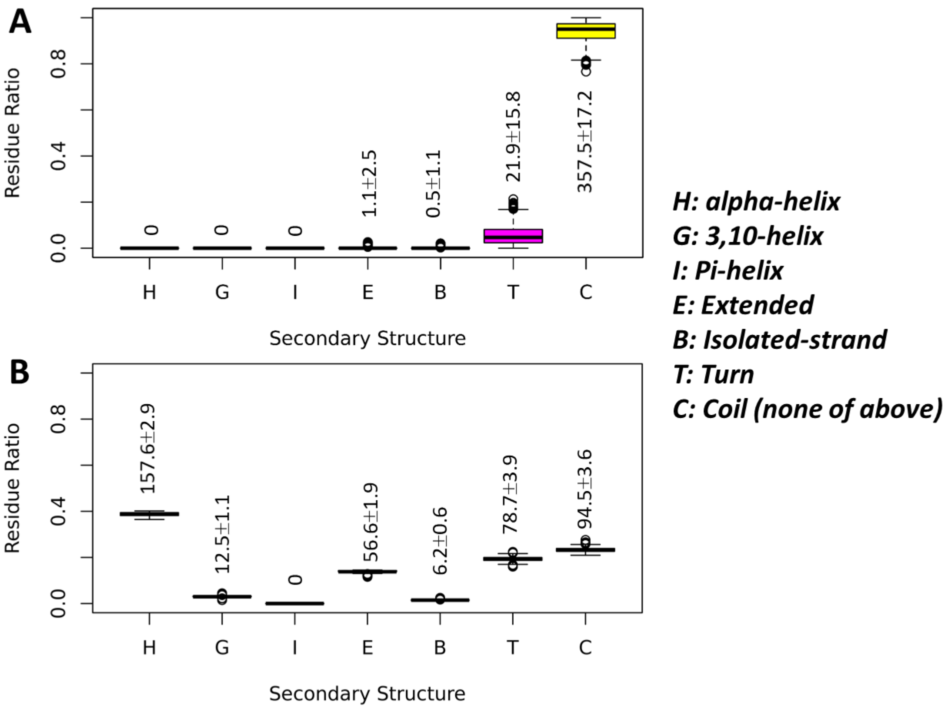


**Figure S5**. Percentages of secondary structure elements (SSEs) estimated in 1000 AF2 models of (**A**) *Nvjp-1* (381 AA) and (**B**) *T7RdhA* (406 AA). The numbers of residues (mean±sd) belonging to each of the seven SSE categories (right side) in the AF2 protein models are given.

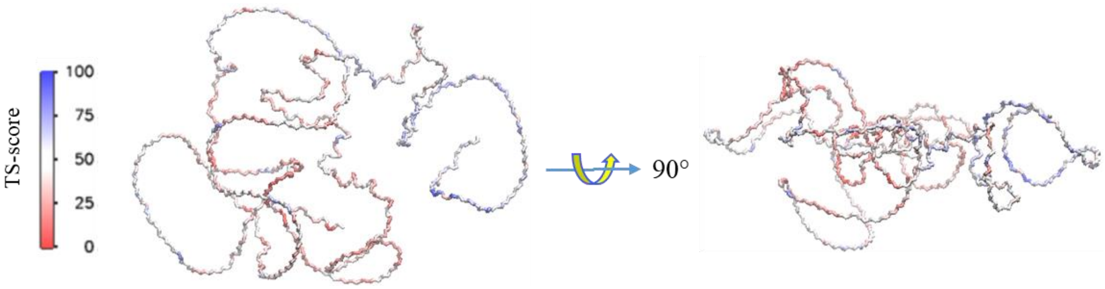


**Figure S6**. The RGN2 model of *Nvjp-1* monomer. All residues are colored by the TS-scores (a measure comparable to pLDDT of AF2) from RGN2. The structure modeling was performed at Google Colaboratory (https://colab.research.google.com/github/aglaboratory/rgn2/blob/master/rgn2_prediction.ipynb).
